# Quorum sensing provides a molecular mechanism for evolution to tune and maintain investment in cooperation

**DOI:** 10.1101/2020.06.29.178467

**Authors:** Eric L. Bruger, Daniel J. Synder, Vaughn S. Cooper, Christopher M. Waters

## Abstract

As selection frequently favors non-cooperating defectors in mixed populations with cooperators, mechanisms that promote cooperation stability clearly exist. One potential mechanism is bacterial cell-to-cell communication, quorum sensing (QS), which can allow cooperators to prevent invasion by defectors. However, the impact of QS on widespread maintenance of cooperation in well-mixed conditions has not been experimentally demonstrated over extended evolutionary timescales. Here, we use wild-type (‘WT’) *Vibrio harveyi* that regulates cooperation with QS and an unconditionally cooperating (‘UC’) mutant to examine the evolutionary origins and subsequent dynamics of novel defectors during a long-term evolution experiment. We found that UC lineages were completely outcompeted by defectors, whereas functioning QS enabled the maintenance of cooperative variants in most WT populations. Sequencing of evolved populations revealed multiple *luxR* mutations that swept the UC lineages. However, the evolution of mutant lineages with reduced levels of bioluminescence (‘dims’) occurred in many WT lineages. These dim variants also decreased other cooperative phenotypes regulated by QS, such as protease production, indicating they result from changes to QS regulation. This diminished investment phenotype optimizes a trade-off between cooperative input and growth output, allowing cooperation to be maintained under QS control even in the presence of evolved defectors.

## INTRODUCTION

Quorum sensing (QS) is a widely distributed form of cell-cell communication used among bacteria [1–4]. It relies upon the production of and response to small signaling molecules called autoinducers that allow cells to alter their patterns of gene expression in response to changes in population density [5–7]. In *Vibrios*, many cellular behaviors have been shown to be regulated by QS, including biofilm formation, secreted products such as surfactants, virulence factors and extracellular enzymes, and bioluminescence [8–16]. Many products that QS regulates are metabolically costly to make and potentially beneficial even to population members that do not contribute to their production, thus falling under the definition of cooperative behaviors. It has been suggested that investment in signaling should be highest in intermediate frequency mixtures of cooperators (producers) and defectors (non-producers), where it would provide maximum information about levels of relatedness within a population [17], while other studies suggest that selection should favor decreased investment in QS [18]. Natural bacterial populations often contain mixtures of cooperators and defectors for public goods and quorum sensing [19–20], raising questions over how much (and in what ways) QS is under selection in natural environments. These observations suggest that selection in nature favors the evolution and persistence of these defecting types, but that these defectors cannot completely sweep natural populations, leading to the coexistence of different cooperative strategies.

We have previously shown that functional QS in mixed populations of *Vibrios* stabilizes cooperative behaviors in the presence of genetically engineered defectors [21], and QS can also promote the increase of cooperative behaviors when competing strains undergo dispersal events or population bottlenecks [22]. These investigations of cooperation and defection in *V. harveyi* utilized staged competitions of mixed populations with the engineered Δ*luxR* mutant that lacks the gene for the QS master regulator necessary to induce high-cell-density genes. Individual Δ*luxR* cells are defectors for positively QS-regulated cooperative behaviors. Engineered deletion mutants of QS master regulators have also been shown to invade as defectors in other experimental systems including *Pseudomonas aeruginosa* and *Vibrio cholerae* [23–27]. Although such studies have been informative in discovering trade-off’s associated with the maintenance of QS, deletion mutations in master regulator genes can also bear significant pleiotropic fitness costs as they often regulate large numbers of genes. For instance, the characterized *luxR* regulon in *V. harveyi* consists of around 10% of the genome [3,28–29]. This architecture means that the QS regulon may include genes relevant to both public and private traits, and this coregulation potentially provides a means to stabilize cooperative behaviors, although this has not been explicitly demonstrated in *Vibrios* [30]. Additionally, although using seeded defectors *a priori* allows tests to see if predictions hold, it does not directly establish the ecological potential for QS defection and test if, in fact, it evolves and how. Nor does it inform on the evolutionary origin and maintenance of the trait (in this case a QS regulation of a cooperative public good), but merely how it contributes in a given environment.

Based on results from short-term selection experiments, we hypothesized that fitter defectors would evolve to potentially invade *V. harveyi* cooperator strains during the course of extended growth in media containing casein as the sole carbon source, an environment in which population yield is dependent on the extent of QS-regulated protease production [21–22]. To examine this hypothesis, we initiated twelve replicate populations of *V. harveyi* cells in well-mixed populations passaged for 2,000 generations from two cooperator genotypes, the WT strain with a functional QS system and the unconditional cooperator (‘UC’) strain that is genetically locked in the high-cell-density QS state leading to constitutive cooperation. We found that although the UC strain was rapidly swept by *de novo* evolved defectors. However, the WT strain reduced defector invasion and maintained cooperation due to the evolution of “dim” mutants that decrease investment in cooperative traits by modulating the QS pathway. These results suggest that QS control of cooperative traits provides a molecular mechanism that provides greater evolutionary potential for cooperators to compete with defectors by enabling bacteria to tune their investment in cooperation.

## MATERIALS AND METHODS

### Evolution Experiments

Six replicate populations of wild-type, Δ*luxOU*, and corresponding *lacZ*-tagged strains of *V. harveyi* (reclassified *V. campbellii* [31]) were passaged in M9 salt medium (Difco) with 2% sodium chloride supplemented with a) casamino acids (Difco) or b) sodium caseinate (Sigma) (referred to as ‘M9-casein’ and ‘M9-CAA’ herein). This produced 48 populations (Table 1). Cultures (1 mL) were passaged in 96 deep-well plates, and diluted daily 1000-fold into identical media (~10 generations/day). Frozen stocks (20% glycerol) were made periodically and stored at −80 °C. Populations were sampled to determine growth, bioluminescence, and protease production. If contamination of cultures was experienced, evolution was reinitiated from the most recent frozen stock available.

**Table 1.** Strains used in this study. Populations analyzed by sequencing are highlighted in bold. WT populations with significant dimming effects observed are marked with an asterisk.

| Founder and Control Strains: |  |  |  |  |  |
| --- | --- | --- | --- | --- | --- |
| Strain | Genotype | Reference |  |  |  |
| BB120<br>(ATCC BAA-1116) | Wild-type<br>*Vibrio campbellii* | [78] |  |  |  |
| ELB714 | Wild-type<br>attTn7::lacZ | This Study |  |  |  |
| JAF78 | $\Delta luxOU$-Cm<sup>R</sup> | [79] | | | |
| ELB738 | $\Delta luxOU$-Cm<sup>R</sup><br>attTn7::lacZ | This Study | | | |
| KM669 | $\Delta luxR$ | [80] | | | |
| Experimental Populations: |  |  |  |  |  |
| Population | Initiating strain | Lac Colony Phenotype | QS phenotype | Media Treatment | Defector Frequency<br>2000 generations |
| WTcas01* | BB120 | White | WT | M9-casein | 58 |
| WTcas02* | ELB714 | Blue | WT | M9-casein | 38.42 |
| WTcas03* | BB120 | White | WT | M9-casein | 0.28 |
| WTcas04* | ELB714 | Blue | WT | M9-casein | 37.46 |
| WTcas05* | BB120 | White | WT | M9-casein | 0.34 |
| WTcas06 | ELB714 | Blue | WT | M9-casein | 80.93 |
| WTcas07 | BB120 | White | WT | M9-casein | 95.95 |
| WTcas08* | ELB714 | Blue | WT | M9-casein | 2.89 |
| WTcas09 | BB120 | White | WT | M9-casein | 100 |
| WTcas10* | ELB714 | Blue | WT | M9-casein | 74.54 |
| WTcas11* | BB120 | White | WT | M9-casein | 55.62 |
| WTcas12 | ELB714 | Blue | WT | M9-casein | 13.09 |
| UCcas13 | JAF78 | White | UC | M9-casein | 100 |
| UCcas14 | ELB738 | Blue | UC | M9-casein | 100 |
| UCcas15 | JAF78 | White | UC | M9-casein | 100 |
| UCcas16 | ELB738 | Blue | UC | M9-casein | 100 |
| UCcas17 | JAF78 | White | UC | M9-casein | 100 |
| UCcas18 | ELB738 | Blue | UC | M9-casein | 100 |
| UCcas19 | JAF78 | White | UC | M9-casein | 100 |
| UCcas20 | ELB738 | Blue | UC | M9-casein | 100 |
| UCcas21 | JAF78 | White | UC | M9-casein | 100 |
| UCcas22 | ELB738 | Blue | UC | M9-casein | 100 |
| UCcas23 | JAF78 | White | UC | M9-casein | 100 |
| UCcas24 | ELB738 | Blue | UC | M9-casein | 100 |
| WTcaa25 | BB120 | White | WT | M9-CAA | 100 |
| WTcaa26 | ELB714 | Blue | WT | M9-CAA | 100 |
| WTcaa27 | BB120 | White | WT | M9-CAA | 41.48 |
| WTcaa28 | ELB714 | Blue | WT | M9-CAA | 99.32 |
| WTcaa29 | BB120 | White | WT | M9-CAA | 100 |
| WTcaa30 | ELB714 | Blue | WT | M9-CAA | 100 |
| WTcaa31 | BB120 | White | WT | M9-CAA | 100 |
| WTcaa32 | ELB714 | Blue | WT | M9-CAA | 100 |
| WTcaa33 | BB120 | White | WT | M9-CAA | 10.53 |
| WTcaa34 | ELB714 | Blue | WT | M9-CAA | 100 |
| WTcaa35 | BB120 | White | WT | M9-CAA | 56.41 |
| WTcaa36 | ELB714 | Blue | WT | M9-CAA | 100 |
| UCcaa37 | JAF78 | White | UC | M9-CAA | 100 |
| UCcaa38 | ELB738 | Blue | UC | M9-CAA | 100 |
| UCcaa39 | JAF78 | White | UC | M9-CAA | 100 |
| UCcaa40 | ELB738 | Blue | UC | M9-CAA | 12.5 |
| UCcaa41 | JAF78 | White | UC | M9-CAA | 100 |
| UCcaa42 | ELB738 | Blue | UC | M9-CAA | 100 |
| UCcaa43 | JAF78 | White | UC | M9-CAA | 88.65 |
| UCcaa44 | ELB738 | Blue | UC | M9-CAA | 100 |
| UCcaa45 | JAF78 | White | UC | M9-CAA | 100 |
| UCcaa46 | ELB738 | Blue | UC | M9-CAA | 100 |
| UCcaa47 | JAF78 | White | UC | M9-CAA | 100 |
| UCcaa48 | ELB738 | Blue | UC | M9-CAA | 100 |

### Cell growth conditions

#### Artificial Induction of QS

When used, HAI-1 (N-(3-hydroxybutyryl)-homoserine lactone, Sigma-Aldrich 74359) and AI-2 (the furanosyl-borate-diester 3A-methyl-5,6-dihydro-furo [2,3-D][1,3,2] dioxa borole-2,2,6,6A tetraol, gift from B. Bassler) were added to 10 μM (final concentration) as previously described [32]. In competition experiments, populations were seeded with 1% Δ*luxR* as portion of the populations and diluted to 1/1000 of carrying capacity in M9-casein to initiate (as in [21]).

### Growth productivity estimates

Growth was measured by diluting sample, plating on LB agar, and enumeration of resulting colonies or by measuring absorbances (600 nm) of population subsamples in a Beckman Coulter spectrophotometer (Model DU730 A23616, 1 cm path length) and in PerkinElmer EnVision or Spectramax M5 plate-readers.

### Phenotype measurements

#### Protease production

Extracellular protease production of clones from populations at generations 870 was determined by FITC-caseinase assays (Sigma-Aldrich PF0100) as previously described [21]. Concentrations were estimated by normalization to a trypsin standard curve. Monocultures were passaged in LB (24 hours), diluted 1000-fold into M9-casein media (24 hours), and then filtered culture supernatants were assayed. Data from evolved variants were fit with a Michaelis-Menten curve, using GraphPad Prism 8.4.1 software.

#### Bioluminescence measurements

Bioluminescence was measured in 100 μL culture (or a 10-fold dilution series) in an Envision Multilabel Plate Reader (PerkinElmer). Clonal bioluminescence was also qualitatively assessed by examining colonies on LB agar in an AlphaImager HP light box (ProteinSimple, chemoluminescence filter, no visible light applied). Colonies were phenotypically classified (bright, dim, defector) based upon visual examination.

#### Defector determination

In Figure 1, Δ*luxOU* and Δ*luxR* descendants were identified by screening for chloramphenicol resistance (LB-chloramphenicol (10 μg/mL)). The Δ*luxOU* strain was distinguished from Δ*luxR* by colony PCR of selected clones.

**Figure 1.**
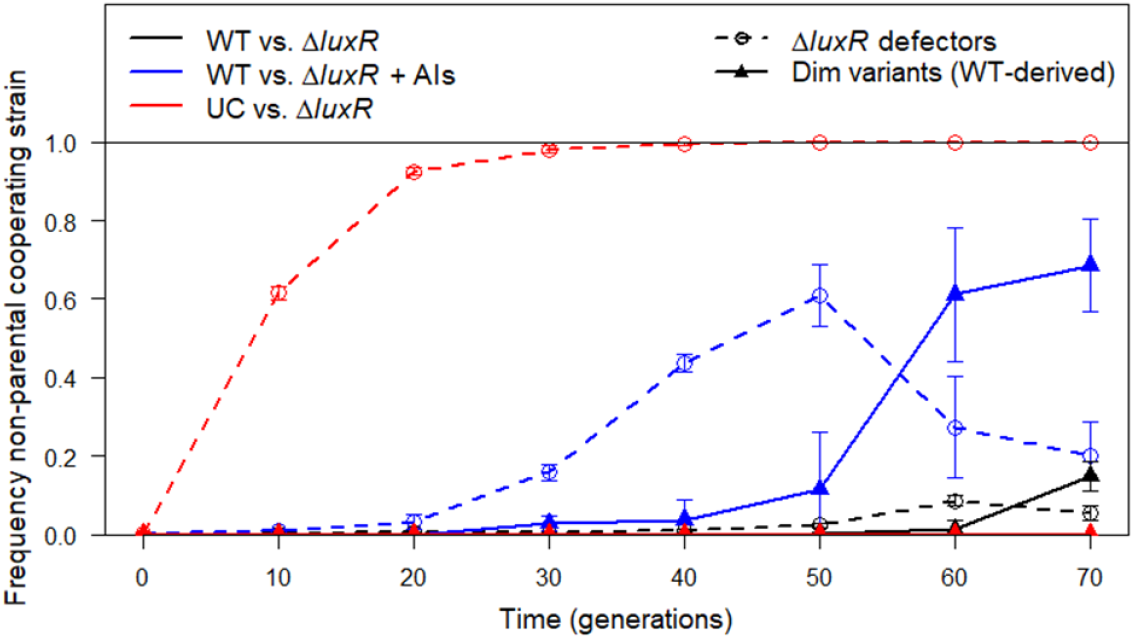
Seventy generation short-term evolution of cooperative populations (99%) seeded with Δ*luxR* defectors (1%). The frequency of defectors or dims when cooperator QS was natively regulated (WT, black), artificially induced by supplementation with exogenous autoinducers (WT + AI, blue), and genetically induced (Δ*luxOU*, UC, red). Two lines are displayed for each treatment group: dashed lines with circles represent the frequency of Δ*luxR* defectors, while solid lines with triangles represent the frequency of evolved dim variants. Error bars represent 95% confidence intervals (N = 5 biological replicates).

### Sequencing and Bioinformatic Analysis

Illumina Nextera barcoding kits were used to generate samples for population sequencing 12 of the WT progenitor populations and 6 of the UC progenitor populations taken at 80, 200, 870, and 2000 generations of evolution. Sequencing was completed on an Illumina NextSeq500 to a minimum depth of [30x coverage]. Reads were filtered for quality and analyzed with Trimmomatic, cutadapt, and seqtk, and mutations were identified using breseq [version 0.27.2] in polymorphism mode [33–36].

## RESULTS

### Artificial induction of QS increases selection for reduced investment in cooperation

Previous studies with *V. harveyi* indicated that a WT functional QS system could significantly inhibit invasion by defector cells during competition whereas an unconditional cooperator (‘UC’, Δ*luxOU*) strain was rapidly invaded [21]. One primary explanation of this result is that premature production of protease by the UC strain at low densities enhanced defector invasion. By extension, we hypothesized that artificial premature induction of the high-density QS state in the WT strain would decrease the ability of WT to prevent defector invasion. The addition of exogenous autoinducer (AI) signal molecules that activate the QS response would then be expected to elicit a constitutive QS phenotype resembling that of the UC mutant. To test this prediction, we used supplementation with 10 μM of HAI-1 and AI-2, the two predominant AIs produced in *V. harveyi* [32,37]. Addition of these AI signal molecules resulted in WT bioluminescence being virtually indistinguishable from UC at all densities during growth, demonstrating that QS was constitutively induced by their addition (data not shown).

We conducted multi-round competition experiments (7 passages, for 70 generations in total), akin to previous studies [21], of WT with the non-luminescent Δ*luxR* defector (at 99% and 1% frequencies, respectively) with and without AI supplementation. Although this experiment was structured to test the interaction between the seeded strains, and particularly to see if the defector could invade from its rare starting frequency, its duration allows the potential for novel genotypic evolution to occur. With AI supplementation, Δ*luxR* defectors did increase in frequency, starting at 20 generations and reaching an average of ~60% of the population by 50 generations (Fig. 1, blue dashed line). However, at 50 generations we began to observe colonies that exhibited reduced but still detectable levels of bioluminescence (“dim” variants, blue solid line). Screening for chloramphenicol resistance allowed us to distinguish between Δ*luxR* (Cm^R^) and WT (Cm^S^), leading us to conclude that the dim variants had evolved from the WT parent (Fig. 1). Ultimately, dims rose to the majority of the WT population, outcompeting the other strains present, including the Δ*luxR* defector. When competed against UC, Δ*luxR* rapidly increased in the frequency to 100% of the population, and no dim variants were observed (Fig. 1, dashed red line).

The invasion of dims in the WT/Δ*luxR* competition at generations 50-70 corresponded with a reduction in population growth yield (Fig. S1A, blue line). This decrease in growth yield is not as extensive as the collapse that occurs when Δ*luxR* invades UC (Fig. S1A, red line). In the WT strain with no AI added, Δ*luxR* defectors did ultimately begin to increase in frequency after 50 generations (Fig. 1, dashed black line), but this increase was rapidly followed by the evolution of WT dims (Fig. 1, solid black line). At the end of this short-term evolution experiment, examination of numerous WT evolved isolates showed a range of bioluminescent phenotypes. We also observed reduced bioluminescence for the evolved bright clones compared to the parental WT strain, suggesting that even population members lacking a qualitatively distinct dim phenotype are under selection for decreased QS investment under the artificially high signal level condition (Fig. S1B). These experiments suggest that a primary strategy to combat defector invasion is modulation of a functional QS signaling network to decrease investment in cooperation without complete loss of this pathway, thereby optimizing the apparent trade-off between metabolic investment into cooperation and resulting growth output.

### Long-term experimental evolution in strains with different levels of QS function results in different population dynamics

The short-term evolution experiment demonstrated that the WT strain could evolve to produce fitter variants that could withstand defector invasion even under artificially induced conditions. We hypothesized that the evolution of such variants could play a critical role in the maintenance of cooperation within experimental populations in conditions where cheating is possible. To better understand how the evolution of decreased investment in cooperation can occur, we initiated a longer-term evolution experiment in twelve replicate populations of each of two cooperator strains: WT that has a functional QS system and the UC strain that constitutively activates the QS high-density regulon. Importantly, no defectors were added at the start of the experiment and must evolve naturally from the parental cooperator strains. Additionally, this experiment was performed in two different environments: M9-casein in which QS-regulated trait of protease production is required for a high growth yield and M9 + casamino acids (M9-CAA) which contains hydrolyzed casein and does not require protease production (and by extension, QS) to reach a high growth yield. These populations were passaged for approximately 10 generations daily for a duration of 2,000 generations in total.

We expected selection for defectors to be stronger in M9-casein media than in M9-CAA because of the higher potential gains in growth conveyed by protease production in this environment [21]. We used bioluminescence of individual isolates as an indicator of QS-mediated cooperation. Defectors were defined as cells that do not produce any visible bioluminescence, and thus QS-regulated cooperative behaviors, while cooperators were defined as cells that maintain some level of bioluminescence, even if it is reduced from that of the parental strain. As expected, defectors evolved rapidly and fixed in all UC M9-casein populations (Fig. 2, cooperators are yellow and defectors are blue), with two interesting population caveats (discussed in the next section). Figure 3A shows images of isolated colonies displaying their bioluminescence signal across all 12 UC M9-casein populations at different generations of the experiments, demonstrating a rapid loss of bioluminescence (defector colonies are shown in blue as they produce no bioluminescence).

**Figure 2.**
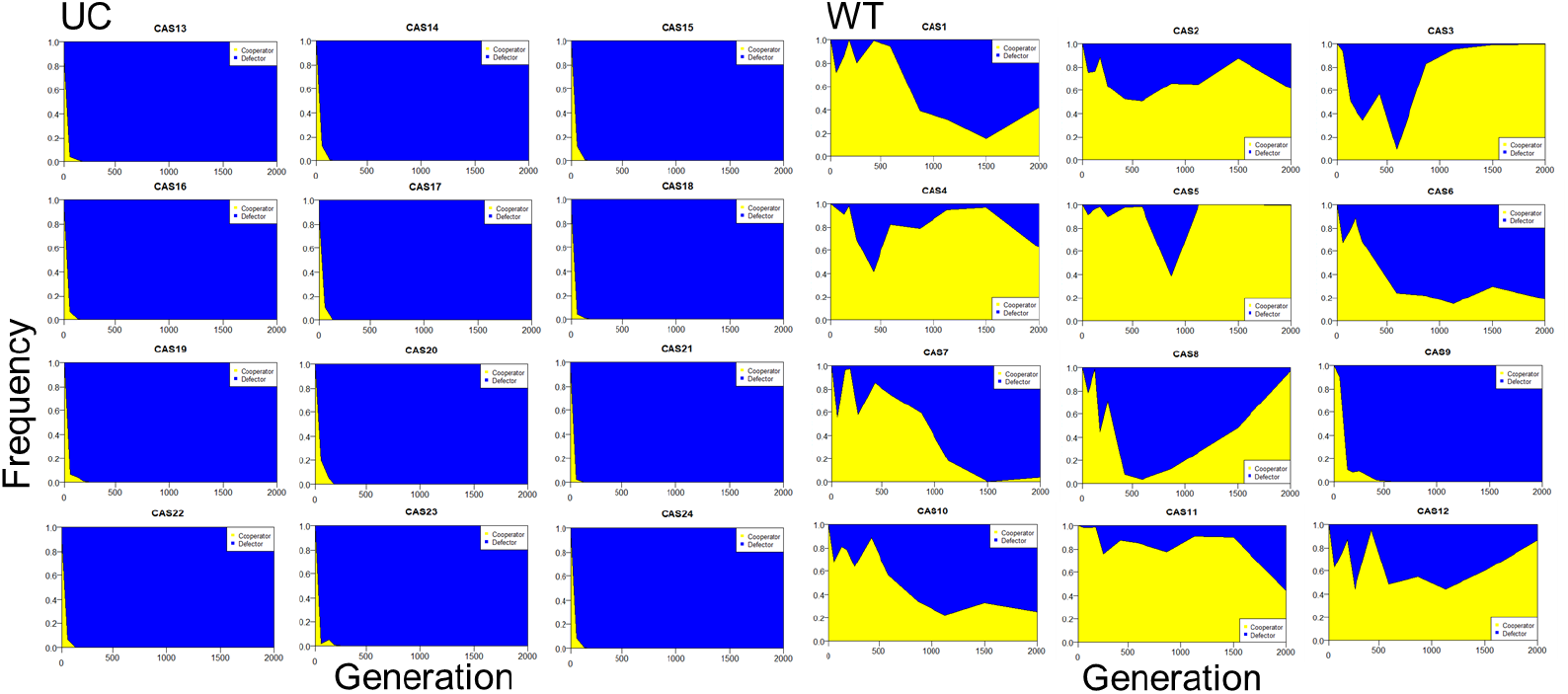
Evolution of cooperative populations in M9-casein medium for 2000 generations. Each panel represents a replicate among the 12 WT and 12 UC lineages. Cumulative frequencies of cooperators (yellow, gauged by visible bioluminescence) and defectors (blue, visibly non-bioluminescent colonies) are tracked over the course of a 2000 generation evolution experiment.

Shortly after the appearance of defectors in the UC lineages, defectors also became detectable in WT M9-casein populations (Fig. 2). However, the rates at which defectors increased in the WT populations was highly varied and they generally did not sweep the WT populations with 11/12 populations maintaining cooperators. Defectors reached an average of 46% across the treatment, although there was high variability in their frequencies between experimental populations (range = <0.01%-100%, Figs. 2, S2A). The maintenance of cooperation in the WT populations was also apparent as most maintained bioluminescent colonies for the duration of the experiment, although the average population-level and frequency of bioluminescent genotypes decreased over time due to the tension provided by selection for defection (Figs. 3B, 4). For the WT lineages, in most cases where defectors are observed, cooperators and defectors coexist together, which has been observed in other experimental systems and conditions [38–42]. Conversely, in the asocial M9-CAA control condition, cooperation was consistently selected against because it was not beneficial in this growth condition (Figs. S2C, S2D). Additionally, the more rapid invasion into UC populations in M9-casein than in M9-CAA was consistent with our prediction of stronger selection for defectors in the casein environment. For the remainder of this study, we will focus specifically on the populations evolved in M9-casein.

**Figure 3.**
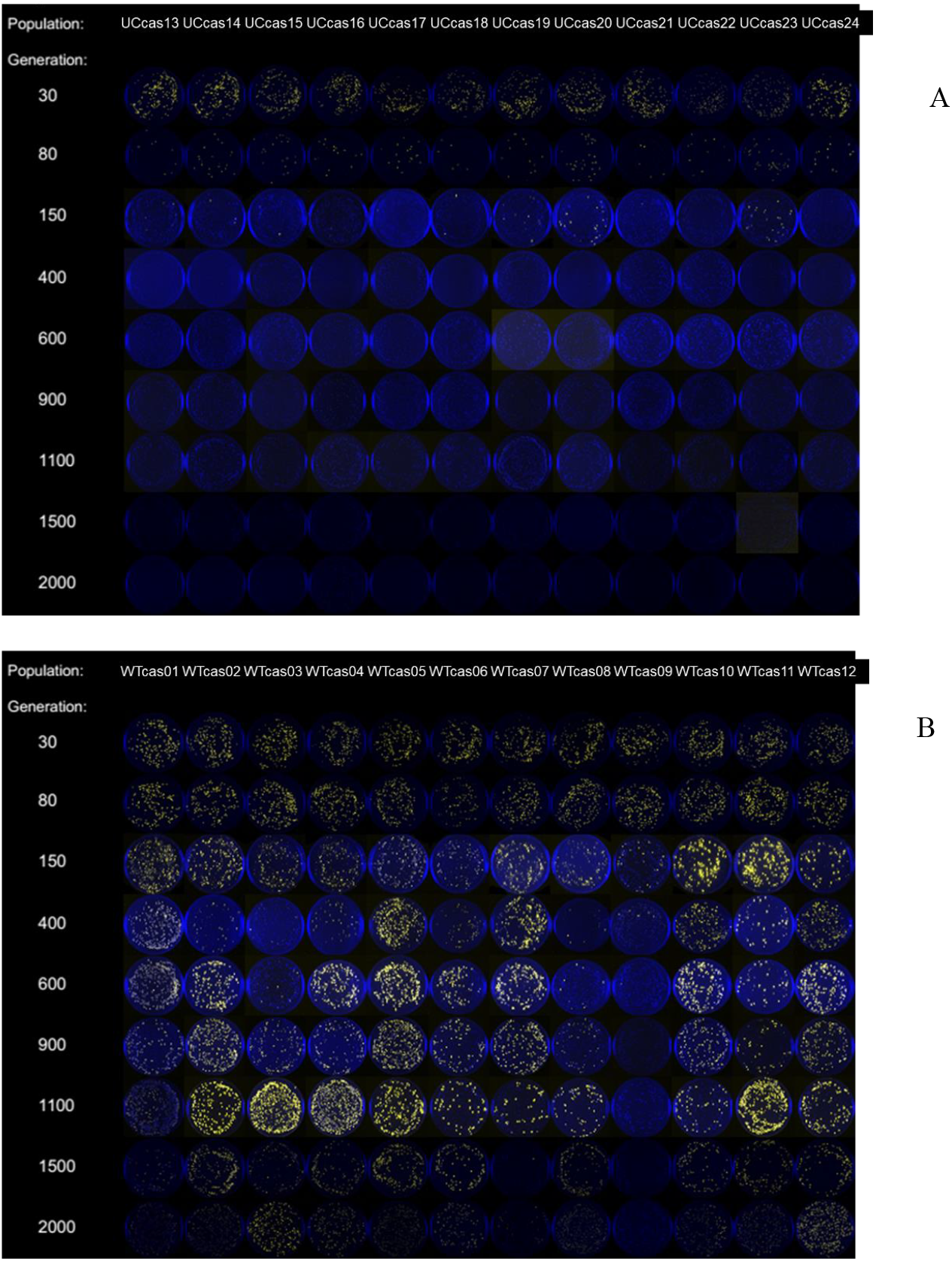
Changes in bioluminescence of M9-casein evolved populations. Over the course of evolution, bioluminescence was monitored by visualizing individuals in the A) UC populations (UCcas13-UCcas24) and B) WT populations (WTcas01-WTcas12). Images are false colored overlays of independent images taken with and without light. Yellow colonies indicate bioluminescent clones and blue colonies indicate non-luminescent clones. Columns indicate separate replicate populations and rows indicate experimental time in generations.

The rise of defectors in populations from both genotypic backgrounds resulted in lower daily population densities once defectors became detectable (Fig. S2B). For UC populations, this was because defectors swept those populations and defection corresponds to decreased casein breakdown and thereby limits nutrient availability. However, drops were also observed in WT populations where defectors predominantly did not sweep. This suggests that cooperators contributed less to public goods in WT populations, either due to the decrease in cooperator frequency or because there was active selection to lessen investment in public goods. The latter is supported by the evolution of dim variants and their increase to sizable portions of their populations.

### Alternative strategists have strong effects on population-level phenotypes

Because the evolved populations were started from clonal WT or UC cooperators, they initially had similarly high levels of bioluminescence (~2.6×10^9^ RLU/mL) and reached high densities on M9-casein (~7.2×10^8^ CFU/mL, Fig. 4). Beyond cooperation alone, this also impacts resulting population biology, as population size has cascading effects on factors including mutational availability and strength of selection. By ~30 generations and then over the course of the experiment, the populations diverged in both properties. At 2000 generations, ten UC populations were phenotypically indistinguishable in both growth (lower yield) and bioluminescence (undetectable) from Δ*luxR* (Fig. 4, solid red circles). By contrast, the other two UC populations (UCcas16, UCcas19) experienced comparable early drops in population density, but rebounded and ultimately experienced large increases in yields, with ending population yields among the highest of all the experimental populations (Fig. S2B). UCcas16 and UCcas19 populations also had detectable but low levels of bioluminescence, which was in some cases only detectable in a sensitive plate reader (Fig. 4). The WT populations display a diverse range of population-level bioluminescence readings at 2000 generations (Fig. 4, solid black circles). Linear regression analysis showed a strong positive relationship between population bioluminescence and maximum density for WT populations (R^2^=0.82, Fig. 4). These results reflect the strong dependence of population-level outcomes, including overall growth upon the types and relative representation of QS phenotype(s) found within the experimental populations; the trend is particularly apparent and continuous among the WT populations.

**Figure 4.**
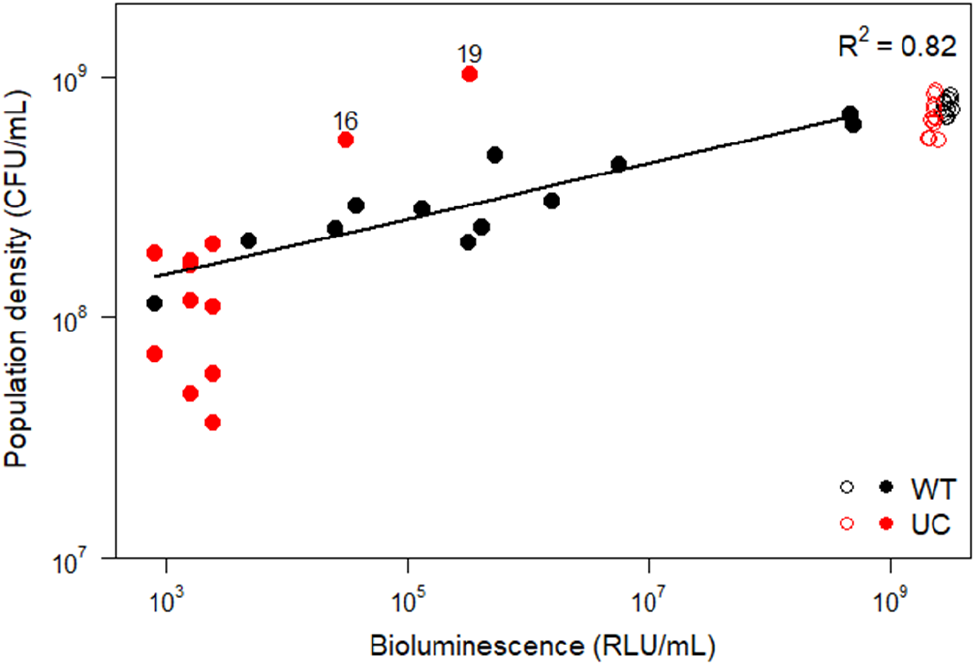
Relationship between population-level bioluminescence (RLU/mL) and population density (CFU/mL) in M9-casein at the initiation of the long-term evolution experiment (open circles) or at generation 2000 (closed circles). WT populations are in black while UC populations lines are in red. Linear regression analysis showed a strong and continuous correlation between these properties for WT populations (R^2^=0.82, p<0.001). The numbers on the graph correspond to the population numbers of the two outlying UC lineages.

### Evolution of *luxR* drives evolutionary dynamics in UC but not WT lineages

We sequenced population samples from temporal transects across the duration our evolution experiment at generations 0, 80, 200, 870, and 2000. This included all 12 WT populations, which displayed a wide range of defector frequencies and population-level phenotypes by the completion of the experiment, as well as six UC lineages, which appeared to have been swept by defectors. We found strikingly different patterns of defector evolution between the two genotypic treatment groups. In particular, all UC populations were rapidly displaced by *luxR* mutant genotypes while no *luxR* mutant alleles exceeded 40% frequency in the WT populations (Figs. 5, S3, Table S1). In fact, only half of the WT populations evolved *luxR* mutations at any generation analyzed, and only rose to intermediate frequencies (5-40%) in those populations. In total, 21 distinct mutations were identified at the *luxR* locus, many of which appeared to be deleterious to its function (i.e., premature stop codon, frameshift, deletion). Of these mutations, 14 were identically observed in both the UC and WT lineages (Table S1). Additionally, in many cases multiple mutant *luxR* alleles occur simultaneously within populations. The *luxR* mutations that occurred early in the experiment in UC populations were ultimately replaced by different variants through clonal interference (Fig. S3; [43–47]). For example, in four UC populations, mutations that fell within the *luxR* CDS were outcompeted by mutations that likely imparted regulatory changes. These putative *luxR* regulatory mutants that had mutations upstream (UCcas14, UCcas16) or downstream (UCcas19, UCcas22) of the gene coding sequence (CDS) outcompeted alternative mutations that fell within in the CDS. In the WT populations where defectors fixed (WTcas09) or nearly fixed (WTcas07), no *luxR* mutations were present at detectable levels.

**Figure 5.**
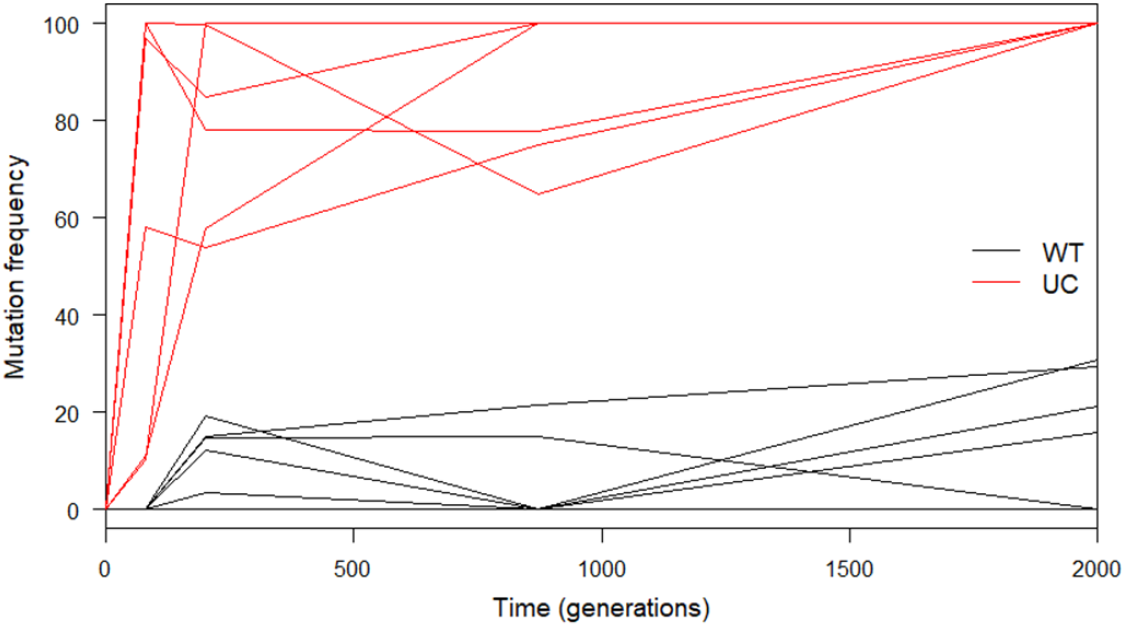
Frequency trajectories of evolved *luxR* mutations in WT lineages where they evolved (black) and UC lineages (red). The line represents the sum of frequencies for all evolved *luxR* variants in a given population.

In contrast to the *luxR* mutations selected in the UC background, 9/12 WT populations exhibited mutations in *luxO*, which encodes a regulatory component of the QS pathway that ultimately represses *luxR* translation at low cell density (Table S2). For the most part (all but WTcas09), the frequencies of these mutants fluctuated over time and did not reach a majority in their population. This finding suggests that a primary target for the evolved dim phenotype is mutated *luxO*. These results demonstrate that although mutation of *luxR* is an effective defector strategy to compete against UC mutants, this strategy cannot sweep WT lineages. Rather, the presence of a functional QS pathway selects for effects like diminished QS activity, (rather than complete loss) of QS by frequently targeting alternative components such as *luxO*.

Additional regulatory targets occurred among both WT and UC populations (Table S2). These included a GntR family transcriptional regulator (VIBHAR_RS03920), the two-component sensor histidine kinase BarA (VIBHAR_RS16525), the two-component system sensor histidine kinase/response regulator VIBHAR_RS23260, the Rsd/AlgQ family anti-sigma factor VIBHAR_RS01060, and an alternative TetR/AcrR family transcriptional regulator VIBHAR_RS19645. The direct impact of all these mutations on QS signaling remains to be determined.

### Evolved dims and defectors possess multiple phenotypic differences from ancestral cooperator strains

As noted, the rise of non-luminescent defectors in evolved populations led to large drops in population densities (Figs. S2B, S2D). Our shorter-term selection experiments showed that evolved dims and non-luminescent defectors can contribute to lower population growth yields (Figs. 1, S1). Because bioluminescence and protease production are induced by QS at high cell density in the ancestor, we wondered whether these traits remained correlated in evolved clones. To test this, evolved clones from the different bioluminescence phenotypic classes from all populations (at generation 870) were measured for bioluminescence and were consistent with their visual classification (data not shown). Protease production was measured when cultures reached their carrying capacity in M9-casein. Evolved bright isolates maintained high levels of protease production while non-luminescent defector clones tested produced low levels of protease analogous to the Δ*luxR* defector (Fig. 6A), confirming that these clones were in fact general QS-defectors. Alternatively, evolved dim variants exhibited intermediate levels of protease production, similar to their bioluminescent phenotype. This result suggests that causative mutations impacted these multiple traits simultaneously rather than separate mutations impacting each process individually.

**Figure 6.**
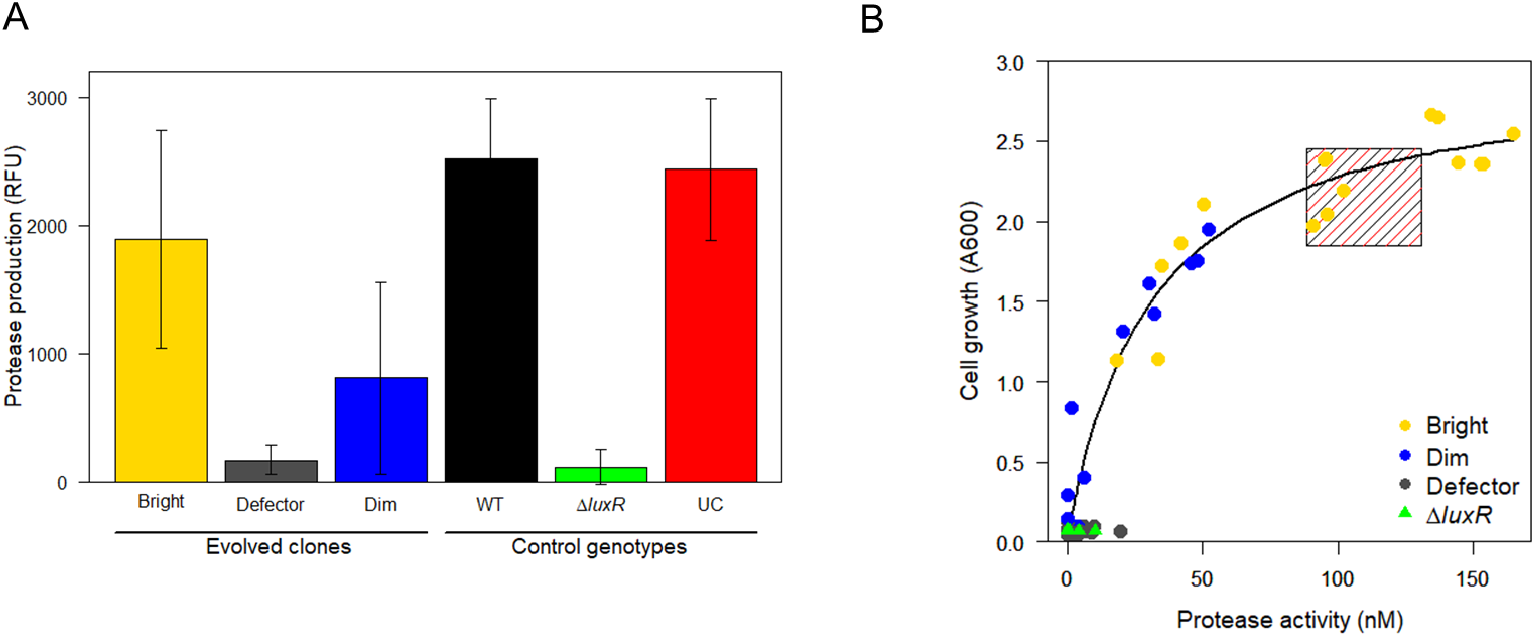
Protease production of evolved clones. Evolved clones that were visually classified as bright (yellow), dim (blue), and dark (gray) were assessed for A) extracellular protease levels by measuring the release of FITC-conjugated to casein molecules and measuring fluorescence. B) Cell growth (measured by Absorbance at 600 nm) was plotted as a function of protease production and fitted with a Michaelis-Menten curve. The WT, Δ*luxR* (green), and UC control strains were replicate cultures initiated from the same genotype, while the evolved clones consisted of multiple genotypes that arose in different populations. The black and red shaded box in panel B represents the 95% confidence intervals for the binned control cooperator strains.

To further understand how varying levels of protease activity impacted growth, we characterized growth phenotypes of these clones in M9-casein when grown in monocultures and compared it to their protease production phenotypes (Fig. 6B). We also measured several replicates of the control genotypes (WT, Δ*luxOU*, Δ*luxR*) for comparison. Among all strains, we found that there was a saturating relationship between the two traits, where clones that produced more protease also grew better in M9-casein media (Michaelis-Menten curve fit, R^2^=0.93). With a Km of 30.85 nM, approximately a third of the amount of protease produced by the founding cooperator strains (mean=106.87-110.70 nM) corresponded to half of the maximum growth yield observed at 24 hours. Beyond this amount, and especially beyond ~50 nM protease (resulting in >80-90% growth of the founding cooperator strains), led to diminishing additional benefits in growth yield. As expected, the Δ*luxR* strain (Fig. 6B, green triangles) displayed relatively low levels of both quantities and the ancestral cooperator strains both possessed high levels of protease production and growth in M9-casein media (Fig. 6B, dashed box). Evolved defector clones had lower levels of growth in the media and produced lower levels of protease, largely resembling the defector control (Fig. 6B, grey circles). Bright variants evolved from the WT parent demonstrated a range of protease production and cell growth that generally resembled the parental cooperators (Fig. 6B, yellow circles) while dim variants evolved from WT all had less protease production and growth than the parental cooperators, although most of these variants were increased relative to the non-luminescent defectors (Fig. 6B, blue circles). Together, these results suggest that evolved WT variants, and especially dims, are optimizing the tradeoff between cooperation and growth in the experimental environment.

## DISCUSSION

Cooperation suffers from an inherent tradeoff between the cost incurred by individuals versus the population-level benefit conferred from performing a particular behavior. As a result, optimizing costs versus benefits provides a challenge to maintaining cooperative behaviors through natural selection. Thus, cooperation is a theoretical challenge for evolutionary biology to explain [48–55]. This study investigated the impacts of QS upon the stability of cooperative behaviors that it regulates over longer evolutionary timescales.

This study also posed the following questions: if defectors were given the opportunity to evolve *de novo*, would they? If so, would the evolutionary paths and mutational targets be comparable between the WT and UC cooperating lineages? To examine the evolution cooperators and defectors, and the resulting impacts on phenotypes, dynamics, and selected genotypes, we incorporated both short-term examination of interactions between cooperator and an engineered defector strain and longer-term evolution experiments. Nearly midway through the evolution experiment (generation 870), a comparison of protease production versus growth for evolved clones revealed a non-linear saturating curve, reminiscent of Monod curves used to describe microbial growth (Fig. 6B, [56]). The coordinates to which many clones evolved is where increases in growth as a function of protease production begin to plateau. This portion of the curve resembles a Pareto front [57–59], suggesting that the evolved WT sublineages with decreased investment in protease production were selected by reducing QS activity. We interpret the dim phenotype as an optimization to the selection regime, where populations that best balance pressures to maximize growth rates and minimize exploitation (namely, maximizing inclusive fitness, [7]), perform well over time.

We predicted that defectors would be less likely to evolve and rise to detectable frequencies in WT populations due to our previous competition results, as well as other reported mechanisms of cheater control such policing and metabolic constraint [15,60–64]. We found that this prediction was partially correct in the conditions tested. Defectors did evolve and increase in the presence of WT, but in significantly reduced proportions compared to the UC ancestor (Fig. 2). All UC populations were rapidly invaded and ultimately swept by evolved variants with dramatically reduced QS activity (Fig. 4), and in nearly all cases were indistinguishable in phenotype from the engineered Δ*luxR* defector strain used as a control. This provides support that QS can act as a form cheater restraint.

We sought to determine the genetic nature of the evolved dims and defectors by sequencing 18 experimental populations. Our shorter-term competition results indicated that UC cannot prevent cheating by Δ*luxR* defectors in the M9-casein environment [21]. It also predicts that the evolved dim variants that invaded WT populations were fitter against the WT ancestor than the Δ*luxR* strain (Fig. 1). Sequencing supported these conclusions as the UC populations were indeed universally swept by *luxR* mutations. Though the exact nature of *luxR* mutations varied between lineages, several were clearly loss-of-function mutations, suggesting decreased cooperation was beneficial (Fig. S3, Table S1). Based on measured frequencies in the WT populations that were swept/nearly swept by defectors, *luxR* mutant alleles could not have accounted for the defector phenotypes present. Because dims outcompeted the Δ*luxR* defectors in the shorter-term evolution experiment (Fig. 1), their intermediate QS phenotypes have a role in preserving cooperation in the populations where they evolve by outcompeting *luxR* null variants. Among the higher frequency *luxR* alleles, many of them were putative regulatory mutations that mapped to the gene’s promoter region, rather than changes to the gene coding sequence [31]. Alternatively, the frequent mutations to *luxO* in WT populations demonstrates that impacting LuxO activity is a beneficial strategy to modulate QS pathways and investment into cooperation without a complete loss of cooperation caused by null mutation in *luxR* (Fig. S3, Table S2).

In the long-term evolution experiment, dim variants were especially enriched for in the WT-casein treatment. Eight of the twelve WT lineages (Table 1) possessed notable population-level dimming or harbored significant dim subpopulations by generation 2000. Notably, WT populations that were swept/nearly swept by defectors by generation 2000 (WTcas07, WTcas09) did not evolve dims. Dim variants were substantially rarer among UC lineages: only two lineages (UCcas16 and UCcas19) displayed any measurable bioluminescence and were so dim that it was difficult to visually detect (Fig. 4). Like true defectors, they exhibited characteristic decreases in population productivity and exhibited no measurable protease activity. However, these populations later regained growth levels and ultimately became among the top performing of all the experimental populations (Fig. S2B, top two red lines). Thus, it seems likely that these extreme dim variants acquired beneficial mutations and became proficient for growth in M9-casein (Figs. S2B, 4). One possibility that we are currently exploring is that protease produced by members of the lineages was a privatized form that remained associated with the producing cell.

The genetic mutations underlying the dim phenotype appear to vary across populations and many regulatory gene targets exist among potential source mutations. Additionally, the observed phenotypic range dims exhibit is quite wide (Fig. 6), suggesting their phenotypes can be conferred by a potentially broad set of mutations and/or gene targets. Evolved mutations include several regulatory genes, suggesting that there are multiple regulatory routes that can impact the degree to which resulting genotypes activate QS (Table S2). Although some of the regulatory targets have not been previously linked to QS in *V. harveyi*, BarA orthologs have been previously described connections to QS signaling in *Vibrio cholerae*, where it has been shown to regulate expression of the QS master regulator HapR, analogous to LuxR in *V. harveyi* [61,65]. Additionally, the GntR homolog in *Serratia* has an observed connection between QS and gluconate metabolism [66]. LuxO can interact with alternative sigma factors (σ^54^ and σ^S^, for example) in *V. harveyi* and other *Vibrio* species [67], suggesting further connections between QS, starvation, and carbon metabolism [68]. Overall, the *luxR* locus does not appear to be the most selectively favored mutational target in WT populations. However, other regulatory genes such those listed above and in Table S2 appear to be common targets. We predict that these targets are more advantageous because they may elicit subtle effects on *luxR* expression, which results in a dim phenotype and maintains cooperation and robust growth.

We observed the evolution of dim variants in M9-casein, where the cost and benefit of cooperative investment is high because this environment requires cooperative protease production for maximum utilization [21]. These evolved dims exhibit reduced cooperation and could potentially outcompete and exploit more extensively cooperating genotypes. However, QS and the regulation of cooperation is maintained in dims, so cooperation in dim-containing populations was preserved. As dims frequently evolved in this study, reduced cooperation appears to be under selection in the experimental conditions. Genotypes with functioning QS were more capable of evolving in response to selective pressure in a robust and repeatable manner. Broadly, the optimization of cooperation seen in dims acts as a mechanism to maintain cooperation even when selection favors defection: the results of this study were observed in well-mixed populations where factors that can favor cooperation, including spatial structural, biofilm formation, and migration did not occur [22,69–70]. This speaks to the robustness of *V. harveyi*’s QS regulatory network, even over long temporal scales in conditions that provide strong selection against cooperation, providing justification for why QS-mediated cooperation is so frequently maintained among microbes. Moreover, we predict that QS promotion of cooperative traits to be even more robust in natural environments where features including spatial structure and migration are commonplace.

Finally, the results of our study provide perspective on the selective forces affecting natural populations where differing degrees of QS are often observed. Particularly, diminished investment in QS has been observed *in situ:* in *Pseudomonas aeruginosa* infections of cystic fibrosis patient lungs, and in diminished infectivity and bioluminescence of natural *Vibrio* isolates [19–20,71–77]. Importantly, we show that functioning QS systems provides robustness against the complete collapse of regulated cooperative behaviors and offer an explanation why mixes of cooperators and defectors, as well as intermediate phenotypes like the dims we describe, are observed in nature.

## Contributions

E.L.B., V.S.C., and C.M.W. designed the experiments

E.L.B. and D.J.S. conducted the experiments

E.L.B. analyzed the data

E.L.B, V.S.C., and C.M.W. wrote/edited the paper

## Acknowledgements

This study was supported by grants R01GM109259 to C.M.W., R01GM110444 to C.M.W. and V.S.C., Frank Peabody and Dr. Marvin Hensley fellowships to E.L.B., and funding from the BEACON Center for the Study of Evolution in Action (NSF Cooperative Agreement DBI-0939454) to E.L.B. and C.M.W. Sequencing was conducted at the Microbial Genome Sequencing center (MiGS) at the University of Pittsburgh. We also thank J. Bazurto (https://orcid.org/0000-0001-9012-2260) and B. Koestler (https://orcid.org/0000-0001-7213-0953) for helpful suggestions and careful review of the manuscript.

---

**Figure S1.**
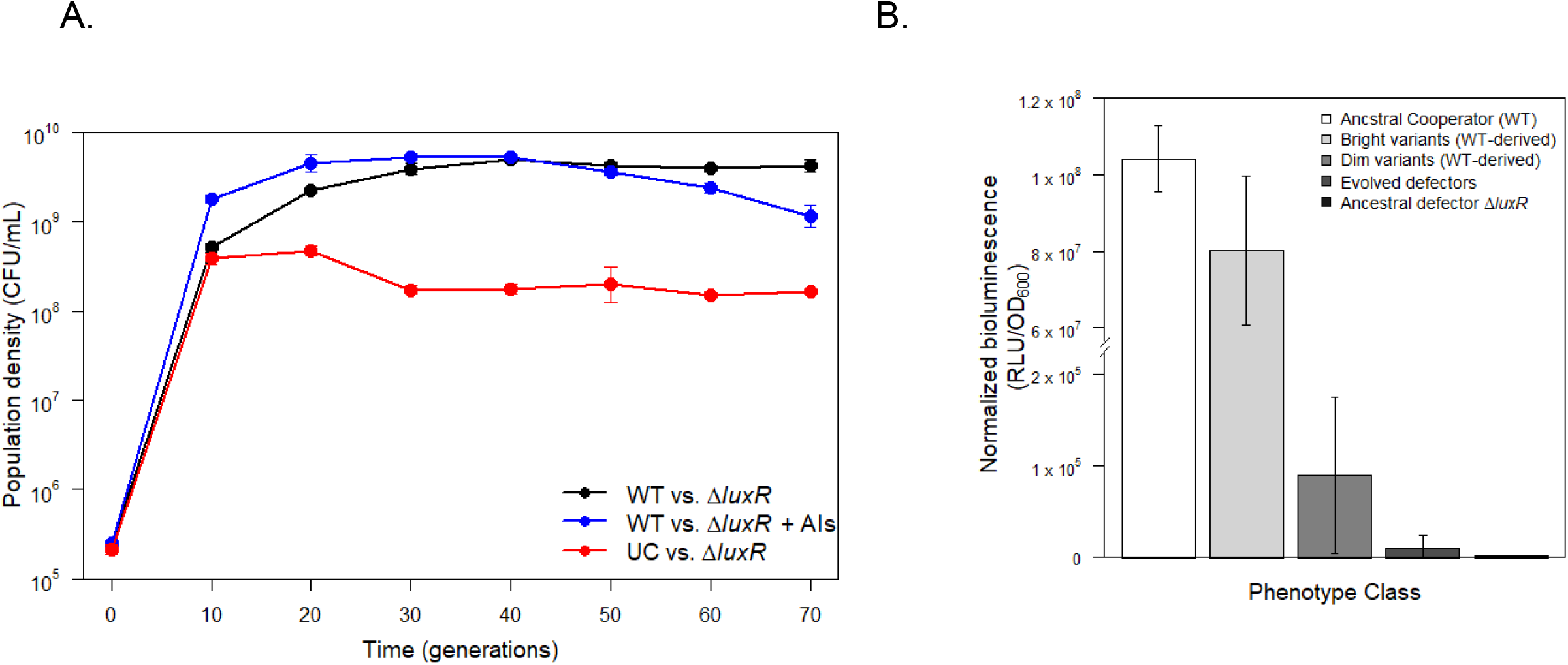
A) Population density during short-term evolution of cooperative populations seeded with Δ*luxR* defectors. Maximum population densities when cooperator QS was natively regulated (WT, black), artificially induced by supplementation with exogenous autoinducers (WT + AI, blue), and genetically induced (Δ*luxOU*, UC, red). Error bars represent 95% confidence intervals (N = 5 biological replicates). B) Comparative outputs for isolates of different phenotypic classes taken at the conclusion of the experiment. These isolates were assessed for bioluminescence as a proxy for QS activity.

**Figure S2.**
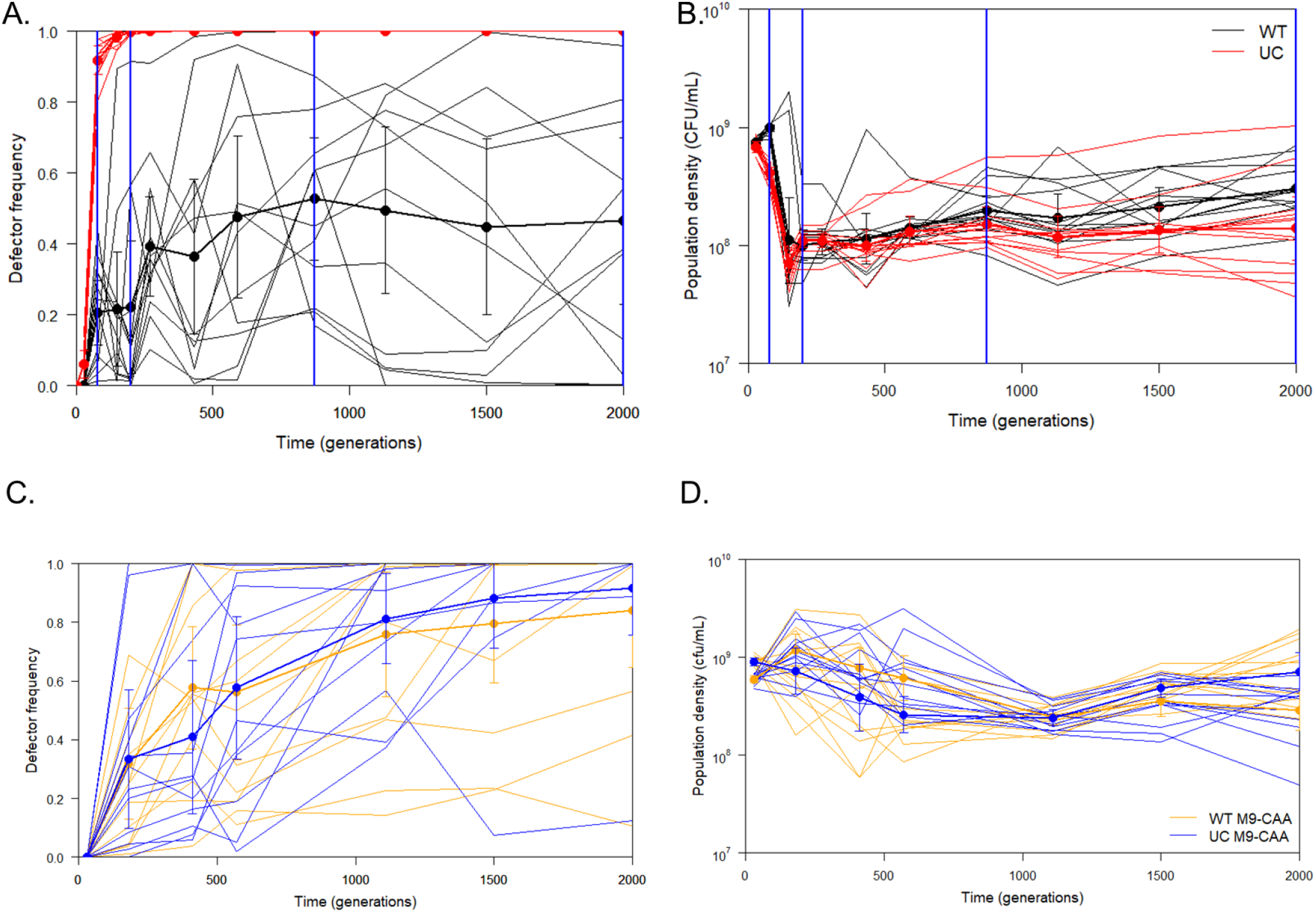
Defector frequency and population density during evolution of cooperative populations for 2000 generations. In each of 12 replicate linages of WT M9-casein (black), UC M9-casein (red), WT-M9CAA (orange), and UC M9-CAA (blue). A and C) frequencies of evolved defectors, and B and D) population densities were quantified over the course of the experiment. C, D) Treatment averages over replicate lineages were determined for more direct comparison to M9-casein evolved populations of WT (black) and the UC (red). In A and B, blue vertical lines correspond to timepoints at which populations were sequenced.

**Figure S3.**
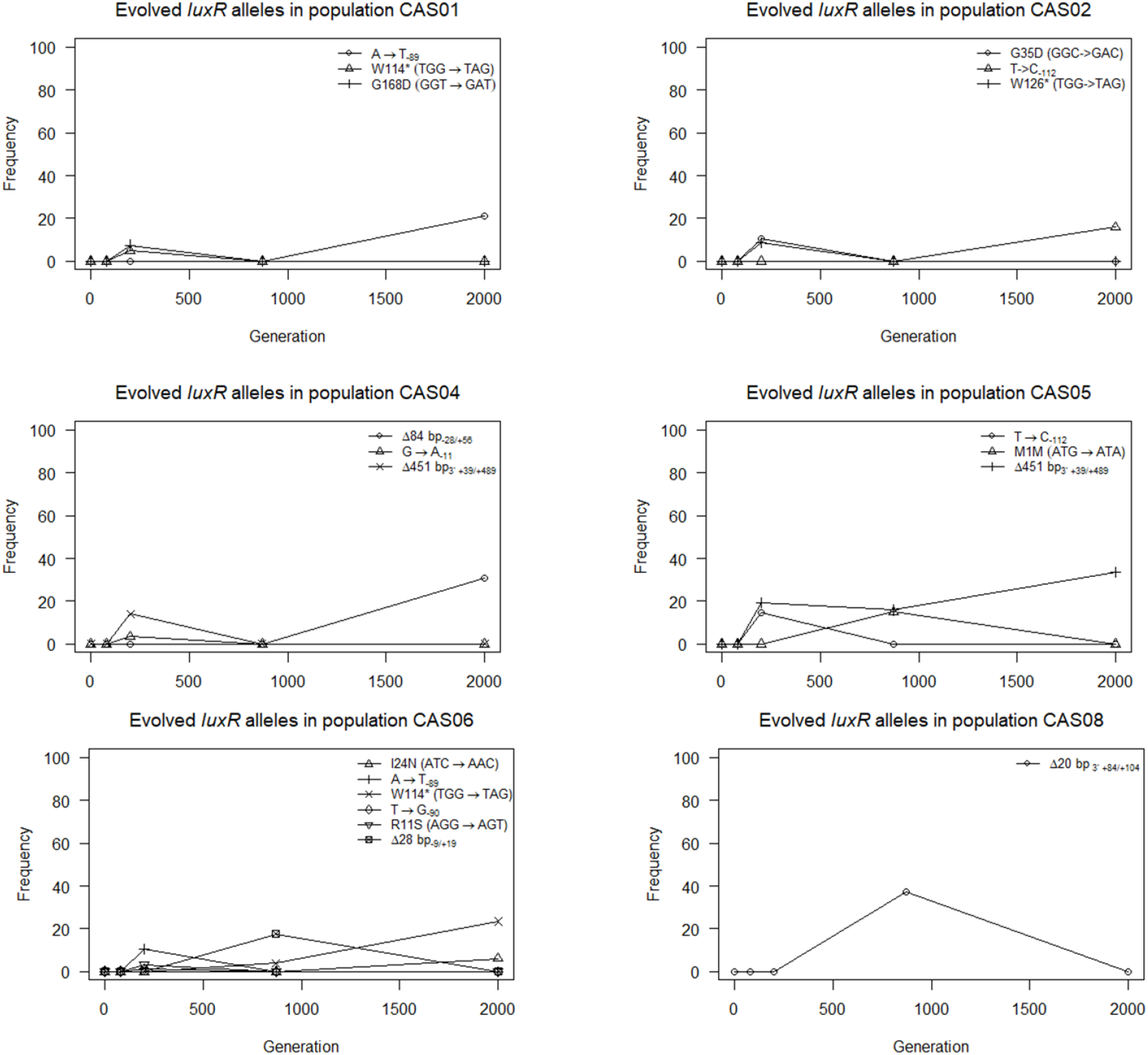
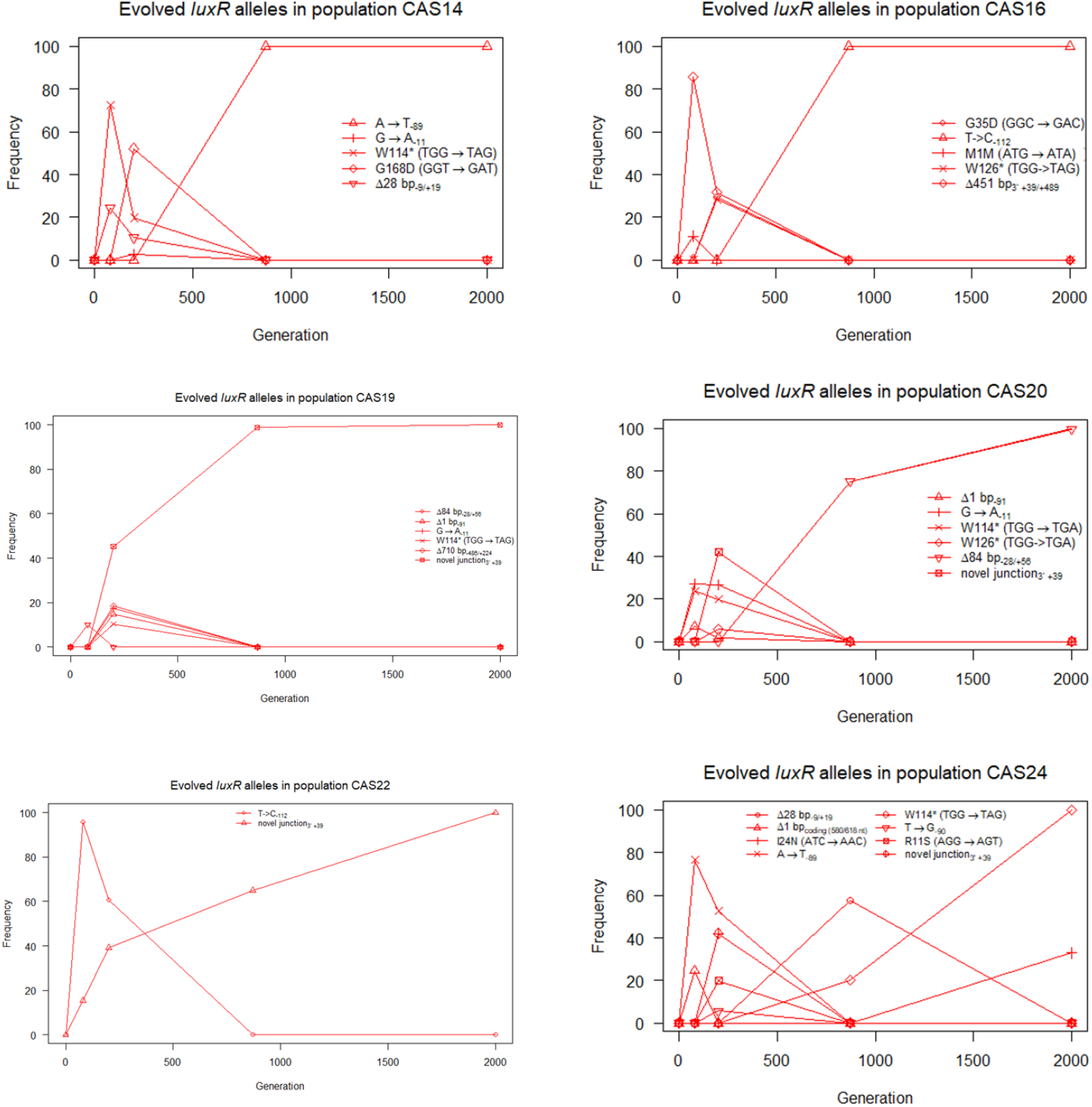
Frequency trajectories of evolved *luxR* mutations divided by population. The abundances of each evolved *luxR* allele in individual WT (black) and UC (red) populations were graphed over time. Each graph contains represents an individual population and details the nature of *luxR* mutations observed within that population.

**Table S1.** Mutations to the *luxR* locus found at detectable levels in experimental populations over 2000 generations of evolution.

| Background strain | Population | Position | Mutation | Annotation | Type | Generation: 0 | 80 | 200 | 870 | 2000 |
| --- | --- | --- | --- | --- | --- | --- | --- | --- | --- | --- |
| WT | WTcas01 | 3,512,670 | G→A | G168D (GGT→GAT) | Nonsynonymous | 0 | 0 | 7.5 | 0 | 0 |
| WT | WTcas01 | 3,512,079 | A→T | intergenic (-233/-89) | Intergenic | 0 | 0 | 0 | 0 | 21.3 |
| WT | WTcas01 | 3,512,508 | G→A | W114* (TGG→TAG) | Truncation | 0 | 0 | 4.8 | 0 | 0 |
| WT | WTcas02 | 3,512,271 | G→A | G35D (GGC→GAC) | Nonsynonymous | 0 | 0 | 10.5 | 0 | 0 |
| WT | WTcas02 | 3,512,056 | T→C | intergenic (-210/-112) | Intergenic | 0 | 0 | 0 | 0 | 15.9 |
| WT | WTcas02 | 3,512,544 | G→A | W126* (TGG→TAG) | Nonsynonymous | 0 | 0 | 8.7 | 0 | 0 |
| WT | WTcas04 | 3,512,140 | Δ84 bp | Deletion (includes start codon) | Deletion | 0 | 0 | 0 | 0 | 30.8 |
| WT | WTcas04 | 3,512,157 | G→A | intergenic (-311/-11) | Intergenic | 0 | 0 | 3.5 | 0 | 0 |
| WT | WTcas04 | 3512823_3759273 | JC | JC_3512823_3759273 | Novel junction | 0 | 0 | 14.07583 | 0 | 0 |
| WT | WTcas05 | 3,512,056 | T→C | intergenic (-210/-112) | Intergenic | 0 | 0 | 14.6 | 0 | 0 |
| WT | WTcas05 | 3512823_3759273 | JC | JC_3512823_3759273 | Novel junction | 0 | 0 | 19.20466 | 15.82382 | 33.55932 |
| WT | WTcas05 | 3,512,170 | G→A | M1M (ATG→ATA) | Synonymous | 0 | 0 | 0 | 15 | 0 |
| WT | WTcas06 | 3,512,159 | Δ28 bp | Deletion (includes start codon) | Deletion | 0 | 0 | 0 | 17.6 | 0 |
| WT | WTcas06 | 3,512,238 | T→A | I24N (ATC→AAC) | Nonsynonymous | 0 | 0 | 0 | 0 | 6.1 |
| WT | WTcas06 | 3,512,078 | T→G | intergenic (-232/-90) | Intergenic | 0 | 0 | 1.3 | 0 | 0 |
| WT | WTcas06 | 3,512,079 | A→T | intergenic (-233/-89) | Intergenic | 0 | 0 | 10.5 | 0 | 0 |
| WT | WTcas06 | 3,512,200 | G→T | R11S (AGG→AGT) | Nonsynonymous | 0 | 0 | 3.1 | 0 | 0 |
| WT | WTcas06 | 3,512,508 | G→A | W114* (TGG→TAG) | Truncation | 0 | 0 | 0 | 4 | 23.4 |
| WT | WTcas08 | 3512868_3512988 | JC | JC_3512868_3512988 | Novel junction | 0 | 0 | 0 | 37.37958 | 0 |
| Δ*luxOU* | UCcas14 | 3,512,159 | Δ28 bp | Deletion (includes start codon) | Deletion | 0 | 24.4 | 10.4 | 0 | 0 |
| Δ*luxOU* | UCcas14 | 3,512,670 | G→A | G168D (GGT→GAT) | Nonsynonymous | 0 | 0 | 52.2 | 0 | 0 |
| Δ*luxOU* | UCcas14 | 3,512,079 | A→T | intergenic (-233/-89) | Intergenic | 0 | 0 | 0 | 100 | 100 |
| Δ*luxOU* | UCcas14 | 3,512,157 | G→A | intergenic (-311/-11) | Intergenic | 0 | 0 | 2.7 | 0 | 0 |
| Δ*luxOU* | UCcas14 | 3,512,508 | G→A | W114* (TGG→TAG) | Truncation | 0 | 72.5 | 19.7 | 0 | 0 |
| Δ*luxOU* | UCcas16 | 3,512,271 | G→A | G35D (GGC→GAC) | Nonsynonymous | 0 | 0 | 29.6 | 0 | 0 |
| Δ*luxOU* | UCcas16 | 3,512,056 | T→C | intergenic (-210/-112) | Intergenic | 0 | 0 | 0 | 100 | 100 |
| Δ*luxOU* | UCcas16 | 3512823_3759273 | JC | JC_3512823_3759273 | Novel junction | 0 | 85.63135 | 31.86559 | 0 | 0 |
| Δ*luxOU* | UCcas16 | 3,512,170 | G→A | M1M (ATG→ATA) | Synonymous | 0 | 11.2 | 0 | 0 | 0 |
| Δ*luxOU* | UCcas16 | 3,512,544 | G→A | W126* (TGG→TAG) | Truncation | 0 | 0 | 28.3 | 0 | 0 |
| *ΔluxOU* | UCcas19 | 3,512,140 | Δ84 bp | Deletion (includes start codon) | Deletion | 0 | 10.1 | 0 | 0 | 0 |
| *ΔluxOU* | UCcas19 | 3,512,077 | Δ1 bp | intergenic (-231/-91) | Intergenic | 0 | 0 | 14.8 | 0 | 0 |
| *ΔluxOU* | UCcas19 | 3,512,157 | G→A | intergenic (-311/-11) | Intergenic | 0 | 0 | 17.5 | 0 | 0 |
| *ΔluxOU* | UCcas19 | 3511681_3512393 | JC | JC_3511681_3512393 | Novel junction | 0 | 0 | 18.42673 | 0 | 0 |
| *ΔluxOU* | UCcas19 | 3512823_3759273 | JC | JC_3512823_3759273 | Novel junction | 0 | 0 | 45.25661 | 98.88741 | 100 |
| *ΔluxOU* | UCcas19 | 3,512,509 | G→A | W114* (TGG→TGA) | Truncation | 0 | 0 | 10.4 | 0 | 0 |
| *ΔluxOU* | UCcas20 | 3,512,140 | Δ84 bp | Deletion (includes start codon) | Deletion | 0 | 0 | 0 | 75.1 | 100 |
| *ΔluxOU* | UCcas20 | 3,512,077 | Δ1 bp | intergenic (-231/-91) | Intergenic | 0 | 7.2 | 1.8 | 0 | 0 |
| *ΔluxOU* | UCcas20 | 3,512,157 | G→A | intergenic (-311/-11) | Intergenic | 0 | 27.2 | 26.6 | 0 | 0 |
| *ΔluxOU* | UCcas20 | 3512823_3759273 | JC | JC_3512823_3759273 | Novel junction | 0 | 0 | 42.03822 | 0 | 0 |
| *ΔluxOU* | UCcas20 | 3,512,509 | G→A | W114* (TGG→TGA) | Truncation | 0 | 23.7 | 19.8 | 0 | 0 |
| *ΔluxOU* | UCcas20 | 3,512,545 | G→A | W126* (TGG→TGA) | Truncation | 0 | 0 | 5.7 | 0 | 0 |
| *ΔluxOU* | UCcas22 | 3,512,056 | T→C | intergenic (-210/-112) | Intergenic | 0 | 95.7 | 60.7 | 0 | 0 |
| *ΔluxOU* | UCcas22 | 3512823_3759273 | JC | JC_3512823_3759273 | Novel junction | 0 | 15.3873 | 39.20275 | 64.9337 | 100 |
| *ΔluxOU* | UCcas24 | 3,512,159 | Δ28 bp | Deletion (includes start codon) | Deletion | 0 | 0 | 0 | 57.6 | 0 |
| *ΔluxOU* | UCcas24 | 3,512,747 | Δ1 bp | coding (580/618 nt) | Deletion | 0 | 24.5 | 0 | 0 | 0 |
| *ΔluxOU* | UCcas24 | 3,512,238 | T→A | I24N (ATC→AAC) | Nonsynonymous | 0 | 0 | 0 | 0 | 32.9 |
| *ΔluxOU* | UCcas24 | 3,512,078 | T→G | intergenic (-232/-90) | Intergenic | 0 | 0 | 5.7 | 0 | 0 |
| *ΔluxOU* | UCcas24 | 3,512,079 | A→T | intergenic (-233/-89) | Intergenic | 0 | 76.6 | 52.5 | 0 | 0 |
| *ΔluxOU* | UCcas24 | 3512823_3759273 | JC | JC_3512823_3759273 | Novel junction | 0 | 0 | 42.03822 | 0 | 0 |
| *ΔluxOU* | UCcas24 | 3,512,200 | G→T | R11S (AGG→AGT) | Nonsynonymous | 0 | 0 | 19.8 | 0 | 0 |
| *ΔluxOU* | UCcas24 | 3,512,508 | G→A | W114* (TGG→TAG) | Truncation | 0 | 0 | 0 | 20.1 | 100 |

**Table S2.**
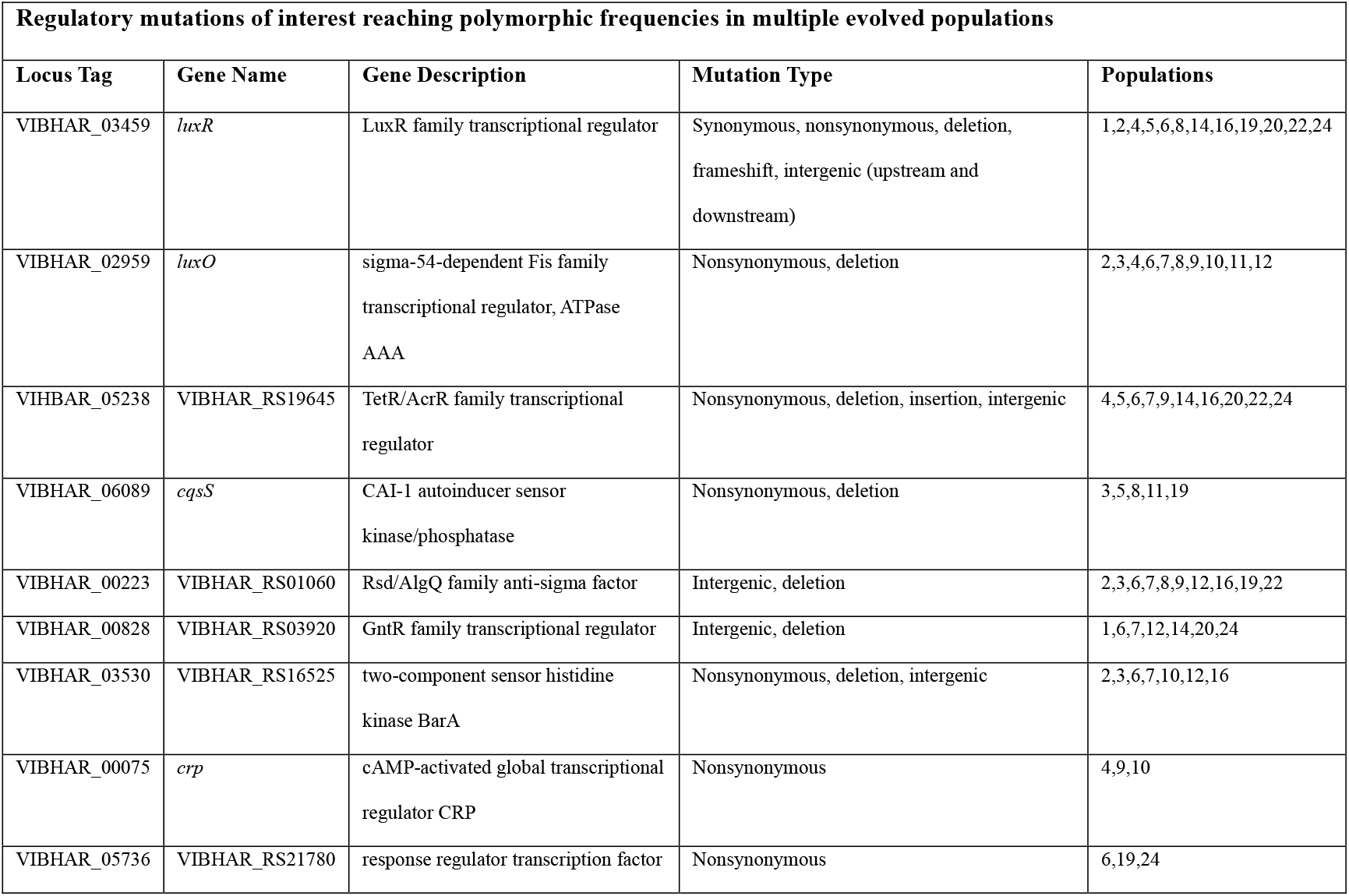
A list of putative regulatory gene hits of relevance to decreasing investment in QS in the experimentally evolved populations.

